# Integrated Metagenomics and Metabolomics Studies Reveal Core Bacterial Guild Regulating Carbohydrate Metabolism in Pediatric MASLD

**DOI:** 10.64898/2026.05.11.724093

**Authors:** Jiating Huang, Xuelian Zhou, Huiying Wang, Ana Liu, Junfen Fu, Guanping Dong, Ying Shen, Wenqing Xiang, Jeffrey Schwimmer, Gang Yu, Jian Huang, Yingping Xiao, Yan Ni

## Abstract

**Background:** Metabolic dysfunction-associated steatotic liver disease (MASLD) is a prevalent pediatric disorder with limited treatment options, primarily due to an incomplete understanding of its molecular drivers. Recent research underscores the role of microbial guilds in metabolic health, but the mechanisms by which dysbiosis driven by core species and co-abundant symbionts disrupt metabolic homeostasis in pediatric MASLD remain unclear.

**Results:** Here, we conducted integrated metagenomic and metabolomic analyses on 285 pediatric subjects including MASLD patients, obese and healthy controls. The gut dysbiosis in MASLD was characterized by a depletion of *Phocaeicola vulgatus*, *Bacteroides uniformis*, *Parabacteroides distasonis*, and *Bacteroides thetaiotaomicron*. Co-abundance network analysis, integrating our cohort with four public datasets, identified these species as core guild members associated with MASLD. Microbial enrichment analysis showed significant disruptions in carbohydrate metabolism, particularly the downregulation of the tricarboxylic acid (TCA) cycle, fructose and sucrose metabolism, and pentose and glucuronate interconversions. *P. vulgatus* and *B. uniformis* were identified as dominant species linked to the downregulation of KEGG orthologs (KOs) in these disrupted pathways that were inversely correlated with hepatic injury biomarkers. CAZyme database analysis further emphasized *P. vulgatus* as the primary contributor to glycoside hydrolases involved in monosaccharide utilization. Finally, both untargeted and targeted metabolomics analysis validated a disrupted metabolic network centered on the TCA cycle and monosaccharide metabolism in pediatric MASLD.

**Conclusion:** Our findings suggest the core guild species *P. vulgatus* and *B. uniformis* may serve as critical regulators of carbohydrate metabolism in pediatric MASLD, offering potential mechanistic targets for gut microbiome-based interventions.

## Background

Metabolic dysfunction-associated steatotic liver disease (MASLD), formerly termed non-alcoholic fatty liver disease (NAFLD), has emerged as the predominant chronic hepatic disorder across pediatric and adult populations [1, 2]. The rebranding to MASLD emphasizes metabolic dysfunction, which carries significant morbidity risks and can progress to cirrhosis and hepatocellular carcinoma, as well as systemic complications such as type 2 diabetes (T2D), cardiovascular diseases (CVD), chronic kidney dysfunction (CKD), and extrahepatic malignancies [3]. Global epidemiological surveillance reveals an escalating prevalence of MASLD, affecting 37.8% of the general population and 3%-12% of children and adolescents. Among overweight and obese youth, the prevalence rises to 39.2% and 52.5%, respectively [4]. However, current therapeutic strategies remain anchored to lifestyle modifications [5, 6], such as dietary optimization and structured physical activity yet clinical outcomes are frequently suboptimal due to poor long-term adherence and incomplete characterization of pediatric-specific molecular drivers.

The gut microbiota, a complex and dynamic ecosystem, relies on core microbial populations and their functions to maintain host physiological balance [7, 8]. Since different microbial taxa do not equally influence ecosystem processes, understanding the ecological roles of community members and the impact of specific taxa on overall biodiversity is crucial. Recent studies have identified core microbiome signatures or keystone taxa that shape community compositions and functions [9]. Additionally, the concept of “guild” [10], drawn from ecology, has recently been applied to gut microbiota research. It refers to groups of bacteria that consistently co-occur and likely cooperate to perform ecological functions [11]. Disruptions in these microbial guild interactions are central to the pathogenesis of various diseases [12, 13]. Wu et *al.*’s research demonstrated that stable but competing microbial guilds within the gut are linked to several systemic disorders, including MASLD-related comorbidities like T2D, CVD, hypertension, and cirrhosis [14]. Compared to the adult MASLD population, fewer research studies have been conducted on the gut microbiome in pediatric MASLD, and most of these studies applied 16S rRNA sequencing approaches with relatively small sample sizes. While the identified microbial features offer initial insights into the gut dysbiosis linked to pediatric MASLD, a more comprehensive understanding of core microbiome signatures and ecological guilds is needed to better capture the structural and functional dysbiosis underlying MASLD pathogenesis.

Moreover, a critical aspect of metabolic regulation lies in the gut microbiota, where microbial enzymes actively participate in the breakdown of nutrients, producing bioactive metabolites that are essential for maintaining metabolic balance. One study estimates that the microbiome explains more than 50% of the variation in circulating metabolites, including gut microbial modifiers and host-microbiota co-substrates [15]. Thus, both metagenomics and metabolomics research have been increasingly applied as a promising strategy for gaining deeper and multidimensional insights into MASLD [16]. Accordingly, advanced bioinformatics tools (i.e., M^2^IA and MetOrigin) have been rapidly developed to facilitate the omics integration analysis and discriminate the microbial metabolites [17, 18]. Early molecular phenomics and metagenomics studies have identified dysregulated molecular networks linking the gut microbiome to metabolic imbalances in aromatic and branched-chain amino acids, which contribute to hepatic steatosis in adult females [19]. Recently, a multi-omics study also found plasma level of N,N,N-trimethyl-5-aminovalerate was positively correlated with the abundance of *Bacteroides* species as an essential feature of MASLD with moderate steatosis [20]. Additionally, growing research has utilized dual-omics strategy to explore the crosstalk between liver-derived bile acids and the gut microbiota in adult MASLD [21, 22]. Despite these advances, the metabolic crosstalk between the host and microbiota in pediatric MASLD remains poorly understood.

To delineate the intricate host-microbiota metabolic interplay in pediatric MASLD pathogenesis, we employed an integrated dual-omics strategy combined with cross-cohort validation: (i) cross-cohort meta-analysis to identify conserved microbial signatures, (ii) cross-cohort co-abundance network analysis to define core microbial interaction modules, (iii) microbial functional exploration using both KEGG and CAZyme databases to identify the most disturbed pathways and modules encoded by specific species, (iv) untargeted and targeted metabolomic profiling to characterize the serum metabolic perturbations. Our findings demonstrated that *Bacteroides species* are the core gut symbionts of pediatric MASLD, and their reduced abundance correlates with disrupted monosaccharide metabolism, potentially contributing to the imbalance of TCA cycle associated with pediatric MASLD development.

## Methods

### Ethical Approval and Study Registration

This study received ethical approval from the Institutional Review Board of the Children’s Hospital of Zhejiang University School of Medicine (Approval Nos. 2021-IRB-038). Written informed consent was obtained from all participants or their legal guardians prior to enrolment. The protocol complied with the Declaration of Helsinki and was registered at the Chinese Clinical Trial Registry (identifier: ChiCTR2100045616; date of registration: 19 April 2021; https://www.chictr.org.cn/showproj.html?proj=63354).

### Participants Recruitment

We enrolled 285 children aged 5-18 years, stratified into three groups based on WHO BMI z-scores [23] and metabolic profiles: (1) Healthy controls (HC, n=55) consisted of normal-weight individuals (BMI z-score ≤ 1.04) without metabolic abnormalities including dysglycemia, hypertension, dyslipidemia followed International Diabetes Federation criteria [24]; (2) Obese controls (OB, n=50) included overweight/obese children (BMI z-score > 1.04/ > 1.64) with no radiological evidence of MASLD (hepatic steatosis < 5% on MRI-PDFF or negative ultrasonography); and (3) MASLD patients (n=180) were overweight/obese individuals with radiologically confirmed hepatic steatosis, defined as MRI proton density fat fraction (MRI-PDFF) ≥5% [25] or ultrasonographic evidence of ≥ Grade 1 hepatic steatosis [26]).

The MASLD group was further subclassified: those meeting criteria for advanced metabolic dysfunction-associated steatohepatitis (MASH) or fibrosis risk (n=64) demonstrated elevated biomarkers (ALT > 80 U/L [26], Fibro-Pen index > 0.28 [27], or liver stiffness measurement (LSM) > 7.1 kPa via FibroScan® PRO [28],, while the simple MASLD subgroup (n=61) maintained lower thresholds (ALT < 60 U/L, Fibro-Pen < 0.05, LSM < 6 kPa).

Individuals were excluded if they met any of the following criteria: (i) metabolically unhealthy normal-weight phenotype; (ii) active or chronic hepatobiliary disorders, including malignancy or viral/autoimmune hepatitis; (iii) neuropsychiatric or systemic conditions that impair the ability to participate; (iv) recent systemic infections or use of hepatotoxic medications/probiotics within the past 3 months and 14 days, respectively.

### Sample and Clinical Information Collection

Prior to sample collection, participants were instructed to maintain a fasting state for ≥12 hours. Blood sampling was conducted through venipuncture during scheduled hospital visits. Fasting venous blood specimens were immediately processed by centrifugation at 1,500×g for 15 minutes at 4°C to isolate serum, followed by aliquoting into cryovials for long-term storage at −80°C. Concurrently, fresh fecal samples were self-collected by participants into sterile, DNA/RNA-free containers, transported to the laboratory under 4°C using ice packs, and frozen at −80°C. Clinical data, including anthropometric indices and biomarker profiles, were retrieved from the hospital’s electronic medical record using standardized case report forms (CRFs) to minimize inter-operator variability.

### Metagenomic Sequencing

Fecal DNA was extracted utilizing the QIAamp DNA Microbiome Kit, with concentration measurements conducted via the Qubit® dsDNA Assay Kit. Library preparation followed the NEBNext® Ultra™ DNA Library Prep Kit protocol, encompassing fragmentation to approximately 350 bp, end-repair, A-tailing, and PCR amplification. The resulting libraries underwent size verification using an Agilent 5400 system (USA) and quantification via qPCR. Paired-end sequencing (2 × 150 bp) was conducted on the Illumina NovaSeq platform (Novogene Co., Ltd., Beijing).

### Sequencing Processing and Taxonomic Profiling

The average sequencing throughput for each sample was around 73.5 million reads. Human-derived reads were removed using BWA-MEM against human reference genome, while adaptors, low quality reads, bases or PCR duplicates were filtered via Trimmomatic (v0.39) as previously described [29]. After quality control and filtering, 65.8 million reads per sample on average were remined and around 89.5% effective reads used in downstream analyses. Taxonomic profiling was conducted with MetaPhlAn4 [30] using default parameters, generating relative abundances across eight taxonomic ranks from phylum to species.

### Co-abundance Network Analysis

Species co-abundance networks were constructed using SparCC [31]. Species included in the analysis were those present in at least 20% of samples. Significant correlations (*P* < 0.05) were filtered for network construction. Network topology index degree was calculated and network visualization was performed in Gephi (v.0.1) with Fruchterman Reingold layout.

### Functional Annotation and Species’ Contribution Calculation

KEGG functional profiling and species’ contributions to community functions was conducted using HUMAnN 3.0 [32]. The analysis began with taxonomic profiling to identify detectable organisms within the samples. Then, Bowtie2 [33] was used to align reads to sample-specific pangenomes, generating alignment files that detailed the mapping of reads to their corresponding gene families. Subsequently, unmapped reads were analysed using DIAMOND’s translated search against the UniRef90 database [34, 35], resulting in additional alignment files for counting hits per gene family. Gene counts from both steps were combined, normalized by read length and alignment quality, and compiled into a comprehensive gene abundance file. Finally, the abundances derived from the UniRef90 search were mapped to Kyoto Encyclopedia of Genes and Genomes (KEGG) Orthology (KO) modules and integrated into structured KEGG pathways.

For CAZyme annotation, paired-end reads were assembled with metaSPAdes using k-mer sizes of 99 to 127 bp [36]. Contigs shorter than 500 bp were filtered out with a custom Perl script. The assembled contigs were aligned using BWA-MEM, and the resulting alignments were indexed with Samtools [37]. CAZyme annotation was conducted using HMMER (v.3.2.1), matching the protein sequences against hidden Markov model libraries from the CAZyme database (v.7) [38, 39]. Species contribution for each CAZyme family was quantified based on matched KO modules from the above HUMAnN analysis.

### Public Dataset Acquisition and Curation

Publicly available whole-genome sequencing (WGS) datasets associated with MASLD were systematically collated from the National Microbiology Data Center (NMDC; https://nmdc.cn) and the NCBI Sequence Read Archive (SRA; detailed information provided in **Table S1**). Inclusion criteria prioritized non-interventional human cohort studies with clearly defined MASLD clinical subgroups (e.g., steatosis, steatohepatitis, fibrosis stages) or healthy controls. Datasets lacking detailed phenotyping, derived from interventional trials, or utilizing non-WGS platforms (e.g., 16S rRNA) were excluded. Ultimately, raw sequencing reads from one pediatric MASLD cohort [40] and three adult cohorts, encompassing MASLD [41], MASH [42], and different fibrosis stages [43], were uniformly reprocessed using standardized pipelines as stated above. Ethical compliance was confirmed for each dataset, adhering to original study approvals and database deposition policies.

### Untargeted Metabolomics Analysis

Serum metabolic profiling was performed using a Waters ACQUITY UPLC system coupled with a Q-Exactive HF-X mass spectrometer (Thermo Scientific) and heated electrospray ionization (HESI-II) in positive/negative switching mode. Four chromatographic methods were applied: HILIC (BEH Amide column, 2.1×150 mm, 1.7 µm; 10 mM ammonium formate, pH 10.8), RP-acidic polar (C18 column, 2.1×100 mm, 1.7 µm; 0.1% formic acid + 0.05% PFPA), RP-acidic non-polar (methanol/acetonitrile gradient with 0.05% PFPA), and RP-basic (6.5 mM ammonium bicarbonate, pH 8). Full-scan MS data (70–1000 m/z) were acquired at 35,000 resolutions, with data-dependent MS/MS (dd-MS²) triggered for top-five ions (dynamic exclusion: 20 s). Raw data were processed using the Discovery HD4 platform (Waters) with LIMS-integrated peak picking, retention time alignment, and compound annotation against an in-house library (>3,300 authenticated standards). Metabolite identification required three criteria: retention index (±2% window), exact mass accuracy (<10 ppm error), and MS/MS spectral match (forward/reverse scores ≥80%). Quality control included internal standard calibration, solvent blank subtraction, and manual artifact removal.

### Targeted Metabolomics Analysis

Serum metabolomic analysis was performed on a Shimadzu LC-30AD UHPLC system coupled to a QTRAP 6500+ mass spectrometer (AB Sciex). Serum aliquots (100 μL) were extracted with methanol/acetonitrile (1:1, v/v) containing isotope-labeled internal standards via vortexing, low-temperature ultrasonication, and protein precipitation (−20°C, 1 h). After centrifugation (14,000 rcf, 4°C), supernatants were filtered through Ostro 96-well plates (Waters). Two analytical strategies were employed: (i) direct analysis of vacuum-dried extracts reconstituted in 50% acetonitrile/water, and (ii) derivatization of aliquots with 3-nitrophenylhydrazine (3-NPH) and EDC prior to reconstitution.

Chromatographic separation used a BEH Amide column (Waters; 2.1 × 150 mm, 1.7 μm) with ammonium formate (10 mM, pH 10.8)/acetonitrile gradient for polar metabolites and a C18 column with methanol/water (0.1% formic acid, 0.05% PFPA) for lipidomics. LC-MS/MS operated in negative ionization mode (−4500 V) using MRM. Quality control samples and retention time calibration standards ensured system stability. Data were processed via Multiquant 3.0.3 (AB Sciex).

### Enrichment Analysis and MetOrigin Analysis

The enrichment ratio of the high class of KEGG pathways and individual KEGG pathways was calculated based on the number of KO with significant abundance (FDR *P* <0.05) in each class and pathway, respectively. Hypergeometric tests were implemented via clusterProfiler package (v4.0) [44] to evaluate whether differentially abundant KOs are statistically overrepresented. Metabolic pathway enrichment analysis was conducted for significantly different metabolites (*P* <0.05) using MetaboAnalyst [45]. The metabolic origins of the detected metabolites were elucidated via MetOrigin 2.0 [17].

### Statistical Analysis

All statistical analyses were performed using R software. Normality of continuous variables was assessed using the Shapiro-Wilk test. For normally distributed data, group differences were assessed by Student’s t-test (two-group) or one-way ANOVA (multi-group). Non-normally distributed variables were analysed using Wilcoxon rank-sum (two-group) or Kruskal-Wallis test (multi-group). Categorical variables were compared via chi-square test. Multiple testing correction via the Benjamini-Hochberg (BH) procedure was applied, adjusted *P* < 0.05 denoted statistical significance.

Bray-Curtis dissimilarity metrics, calculated with the *vegdist* function from the vegan package, were used to assess overall variability in taxonomic, functional, and metabolomic features. Variance explained by MASLD, biomarkers, and covariates was quantified using PERMANOVA with the *adonis* function.

Spearman’s correlation, computed using the *rcorr* function from the Hmisc package, was used to explore associations between clinical indices and differentially abundant species profiles, KO functional profiles, and metabolite profiles. The association between feature prioritization for MASLD phenotypes (e.g., ALT, Fibro-Pen, LSM) and these profiles was further explored using general linear regression models, adjusting for covariates such as age, gender, and BMI z-score.

### Sensitivity Analysis

To address group size imbalances, we performed 100 bootstrapped resamplings of the MASLD group (n = 55 per iteration, with replacement) and applied a hierarchical analytical workflow as described in the section of *Statistical Analysis*. The number of statistically significant comparisons across the 100 resamplings was recorded, and the proportion of significant results was calculated to assess the consistency and stability of the findings.

## Results

### Basic Characteristics of Participants

This study integrated faecal metagenomics and serum metabolomics to delineate core host-microbiota perturbations in pediatric MASLD (**Figure 1A**). A total of 285 children (188 males and 97 females) were enrolled, including HC (33/22), OB (33/17), and MASLD patients (122/58), with a balanced sex distribution across groups (*P* = 0.359, **Table S2**). MASLD participants were older, had higher BMI z-scores, larger waist circumferences, and elevated systolic/diastolic blood pressure compared to both OB and HC groups (all *P* < 0.001). Hepatic injury markers such as ALT, AST, liver stiffness (LSM) and Fibro-Pen indices were significantly elevated in MASLD compared to OB and HC (*P* < 0.05). Dyslipidaemia gradients were evident across the disease spectrum: triglycerides (TG) and low-density lipoprotein cholesterol (LDL-C) progressively increased from HC to OB to MASLD (*P* < 0.001). MASLD patients also exhibited exacerbated fasting hyperglycaemia and elevated HbA1c levels (*P* < 0.05), suggesting systemic metabolic derangement.

**Figure 1.**
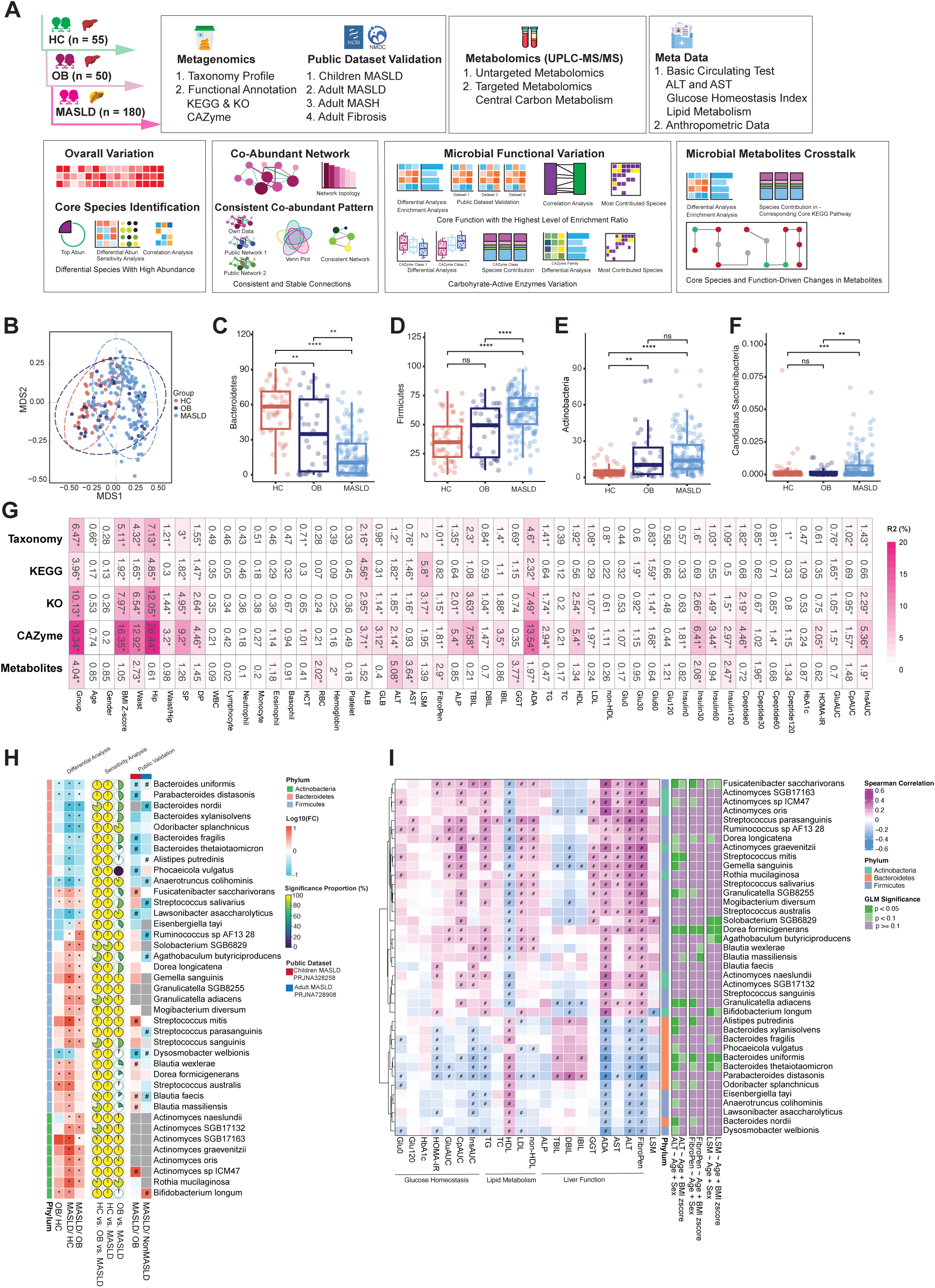
Microbial Signatures with High Abundance in pediatric MASLD. **(A)** Study design. **(B)** Multidimensional scaling analysis plot of taxonomic variation at species level. **(C-F)** Alteration of gut microbiota at the phylum level. Significant changes in the relative abundance of (C) Bacteroides, (D) Firmicutes, (E) Actinobacteria, (F) Candidatus Saccharibacteria. **(G)** Overall variation of omics datasets evaluated by PERMANOVA. The proportion of variation (R2 %) in the profiles of taxonomy, KEGG, KO, CAZyme, and metabolites explained by disease status, covariates, and circulating biomarkers, as quantified by PERMANOVA (with 999 permutations) based on Bray-Curtis dissimilarity. All statistical tests were two-sided. \**P* < 0.05. The abbreviations of the biomarkers are defined in Table S1. **(H)** The log transformed fold change value of microbial species with significant variation (left panel), sensitivity analysis (middle panel), and validation of these changes using public datasets (right panel). Species were compared using the Kruskal-Wallis test across the three groups, with species having FDR-adjusted *P* < 0.05 selected for further pairwise comparison using the Wilcoxon rank sum test (left panel). *FDR *P* < 0.05. The proportion of significant results across the 100 resamplings of MASLD compared with HC and OB respectively were showed in sensitivity analysis (middle panel). The validation of significantly different species across public datasets (right panel). Wilcoxon rank sum test, #*P* < 0.05. **(I)** Correlation between significantly abundant species and clinical indices. Significant correlations (*P* < 0.05, Spearman’s correlation) are represented in color: red indicates positive correlation, while blue indicates negative correlation. The correlations between species and key clinical indices, including ALT, Fibro-Pen, and LSM, were further adjusted for confounders using generalized linear regression. After adjustment, correlations with *P* < 0.05 are shown in deep green, *P* < 0.1 in light green, and *P* ≥ 0.1 in purple.

### Gut Microbiota Perturbations in Pediatric MASLD

We first evaluated the overall microbiome configuration in pediatric MASLD. A clear segregation of gut microbiota was observed from HC to OB to MASLD (**Figure 1B**), primarily driven by a decrease in *Bacteroides* and an increase in *Firmicutes*, *Actinobacteria*, and *Candidatus Saccharibacteria* at the phylum level, especially a trade-off between *Bacteroidetes* versus *Firmicutes* (**Figure 1C-F**, *P* < 0.05). Global PERMANOVA analysis revealed significant MASLD-associated variations in taxonomic composition (R² = 6.47%, *P* < 0.05), microbial KEGG pathways (R² = 3.96%, *P* < 0.05), carbohydrate-active enzymes (CAZyme, R² = 18.34%, *P* < 0.05), and serum metabolites (R² = 4.04%, *P* < 0.05) (**Figure 1G**). Anthropometric parameters, such as hip circumference and BMI z-score, most strongly influenced microbiota structure (R² = 7.13% and 5.11% respectively, *P* < 0.05) and CAZyme variation (16.35% and 26.44% respectively, *P* < 0.05), while ALT predominantly drove metabolite divergence (R² = 5.1%, *P* < 0.05).

At the genus level, we identified that *Bifidobacterium, Bacteroides,* and *Phocaeicola* are dominant genera in the human gut, among which, both *Bacteroides* and *Phocaeicola* were ranking among the top 10 by abundance across public pediatric and adult MASLD cohorts (**Figure S1A**). And these dominant genera, *Bacteroides* and *Phocaeicola* were progressively decreased from HC to OB to MASLD (FDR *P* < 0.05, **Figure S1B**, left panel). Additionally, *Alistipes* and *Parabacteroides* were progressively reduced, while *Blautia* and *Dorea* showed increasing abundance. Sensitivity analysis confirmed the robustness of these changes in *Bacteroides*, *Parabacteroides*, and *Blautia* signatures, with reproducibility greater than 50% across 100 times iterations (**Figure S1B**, middle panel). Moreover, consistent variations of these genera were also found in UK pediatric cohort (PRJNA328258) (**Figure S1B**, right panel).

At the species level, we found *Faecalibacterium prausnitzii*, *Blautia wexlerae* and *Phocaeicola vulgatus* are among the dominant microbial species, and *Phocaeicola vulgatus* was the only one ranking among the top 10 by abundance across both our study and four independent public cohorts (**Figure S1C**). Statistically, a total of 39 differentially abundant species were identified among three groups (FDR *P* < 0.05). Specifically, *Bacteroides uniformis*, *Parabacteroides distasonis*, *Bacteroides fragilis*, *Bacteroides thetaiotaomicron*, and *P. vulgatus* were reduced, while *B. wexlerae*, *Fusicatenibacter saccharivorans*, and *Streptococcus mitis* were enriched in MASLD compared to HC (**Figure 1H**, right panel). These findings were confirmed with 100% reproducibility across sensitivity analysis iterations (**Figure 1H**, middle panel). Moreover, these variations were also replicated with the UK pediatric cohort (*P* < 0.05, PRJNA328258), particularly the reduction of *B. uniformis* was also consistently observed in adult MASLD (*P* < 0.05, PRJNA728908) (**Figure 1H**, right panel).

Correlation analysis with clinical biomarkers revealed significant associations between *B. uniformis*, *P. distasonis*, *B. thetaiotaomicron*, and *P. vulgatus* with hepatic injury/fibrosis markers (Fibro-Pen, ALT), as well as insulin resistance (HOMA-IR). Notably, *B. uniformis* and *B. thetaiotaomicron* maintained statistically significant correlations with ALT after adjusting for age and sex in general linear models (**Figure 1I**). Taken together, the reduction of *B. uniformis*, *P. vulgatus*, *P. distasonis*, and *B. thetaiotaomicron*, alongside the enrichment of *B. wexlerae* and *F. saccharivorans*, may represent core microbial signatures of gut dysbiosis in pediatric MASLD.

### Co-occurrence network analysis shows *B. uniformis, P. vulgatus, and P. distasonis* as core species in MASLD

Co-occurrence network analysis revealed a progressive dysbiosis from HC to OB to MASLD, and *B. uniformis* emerged as a central hub with a high degree of connectivity across all groups (**Figures 2A-C**). In addition, the network exhibited expanded perturbations in MASLD, showing a significantly higher degree (*P* < 0.05, **Figure 2D**) but weaker correlations (*P* < 0.05, **Figure 2E**). Most connections in these three networks were related to the genus *Bacteroides* (**Figure 2F**). The number of connections linked to *Bacteroides* in OB and MASLD was more than twice that in HC, primarily due to the positive correlations between *P. vulgatus*, *P. distasonis*, and other *Bacteroides* species (**Figures 2G-I**). Additionally, in the *Bacteroides*-associated subnetwork of MASLD, *Bacteroides* spp. exhibited more negative correlations with potential opportunistic pathobionts *F. saccharivorans* [40, 46] and *Blautia wexlerae* [47] (**Figure 2I**), underscoring the critical role of *Bacteroides* spp., especially *B. uniformis*, in the gut dysbiosis in MASLD.

**Figure 2.**
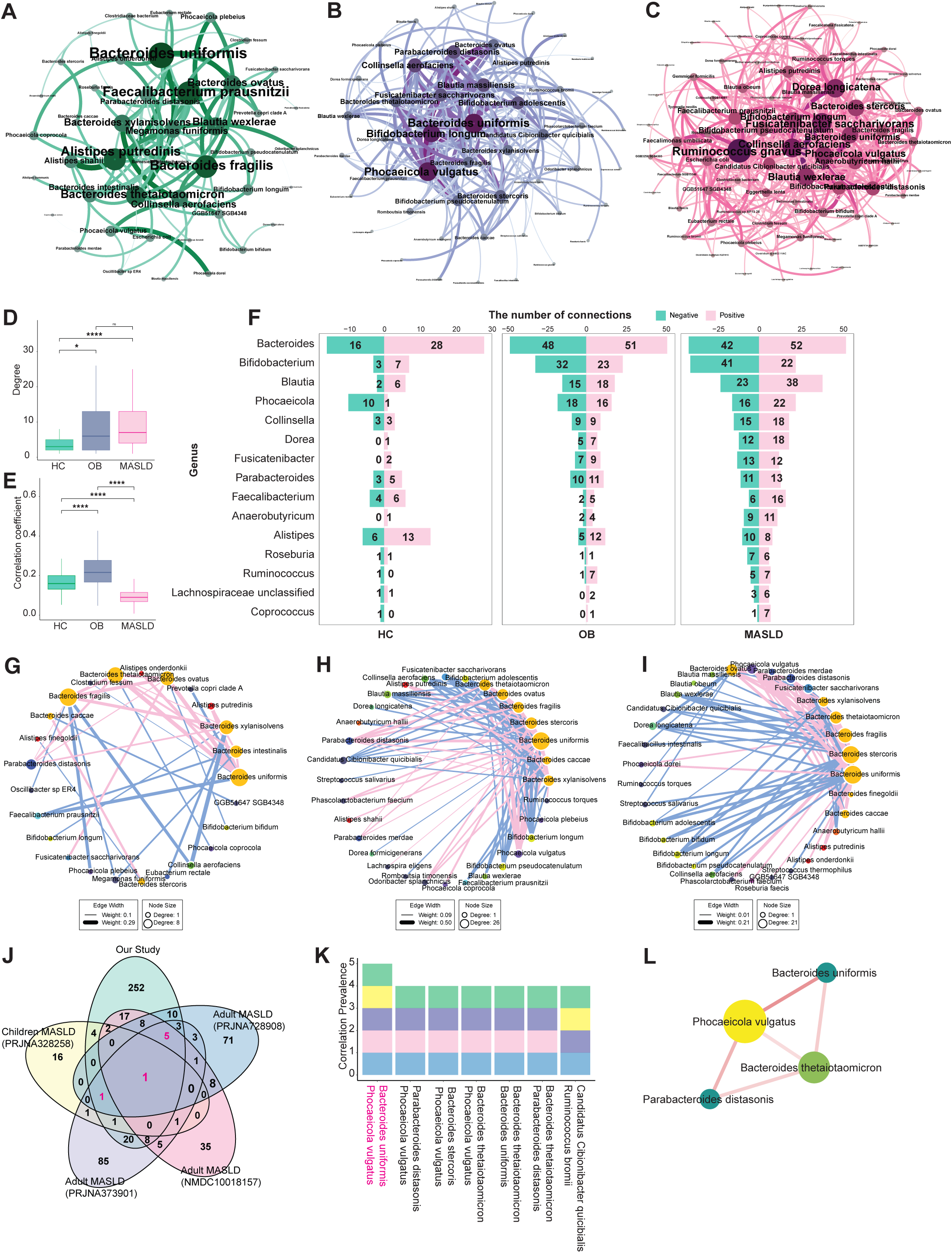
Identification of Core Dysbiosis in Co-Abundant Species of MASLD Patients. (A-C) Co-abundant species networks in HC (A), OB (B), and MASLD (C) individuals, respectively. The color of the lines gradually reflects the absolute value of the correlation coefficient. The size of each node is proportional to its degree. **(D-E)** Comparison of node degrees (D) and correlation coefficients (E) between the HC, OB, and MASLD networks. \*\*\*\**P* < 0.0001, \**P* < 0.05, ns denoted no significance. **(F)** Distribution of interactions for each genus in the microbial networks of HC, OB, and MASLD. **(G-I)** Subnetworks associated with Bacteroides in HC (G), OB (H), and MASLD. Pink lines represent positive correlations, blue lines represent negative correlations, and node size reflects degree. Species from different genera are colored differently. (**J**) Venn diagram showing the number of the shared connections across MASLD-associated networks. **(K)** Correlation prevalence across datasets, with only correlations present in at least four datasets shown. **(L)** Consistent correlations network. Connections consistently present in at least four datasets were selected from our data to construct the network. The red lines represent positive correlations, and the size of the nodes is proportional to their degree.

To confirm the conservation and consistency of co-abundant species signatures in MASLD, we validated the MASLD network using public datasets from both children and adults with MASLD. Consistent co-regulation was observed, with significantly positive correlations between *B. uniformis* and *P. vulgatus* (mean Spearman’s ρ = 0.230 ± 0.074, *P* < 0.05; **Figures 2J-L** and **Table S3**). Additionally, *P. vulgatus* emerged as a keystone taxon, forming significant associations with commensals including *P. distasonis* (mean Spearman’s ρ = 0.236 ± 0.056, P < 0.05), and *B. thetaiotaomicron* (mean Spearman’s ρ = 0.184 ± 0.069, P < 0.05) in our pediatric MASLD network and external networks from three adult MASLD cohorts (**Figures 2J-L** and **Table S3**). These findings highlight *B. uniformis*, *P. vulgatus*, and *P. distasonis* as stable core taxa driving MASLD-associated dysbiosis. Their depletion could destabilize commensal networks, which might facilitate the colonization of pathobionts.

### Gut dysbiosis in pediatric MASLD contributes to the significant disruptions in carbohydrate metabolism

To explore functional changes associated with gut dysbiosis, we identified a total of 45 differentially abundant pathways among three groups (FDR *P* < 0.05), encompassing 326 KEGG orthologs (KOs). The enrichment analysis revealed that carbohydrate metabolism was the most dysregulated functional category (enrichment ratio (ER) = 648.4, FDR *P* < 0.05; **Figure 3A**), with the highest ratio of significantly different KOs to total KOs in this category. Significant changes were observed in both downregulated pathways, such as the tricarboxylic acid cycle (TCA cycle), fructose and sucrose metabolism, and pentose and glucuronate interconversions, as well as upregulated pathways like starch and sucrose metabolism. Additionally, pathways related to glycan biosynthesis and metabolism, including glycosaminoglycan degradation, other glycan degradation, and lipopolysaccharide biosynthesis, were significantly suppressed in MASLD (**Figure 3A**). Notably, carbohydrate metabolism and glycan metabolism accounted for the majority of pathways with the highest enrichment ratios, indicating the most significant functional changes in KOs within these categories.

**Figure 3.**
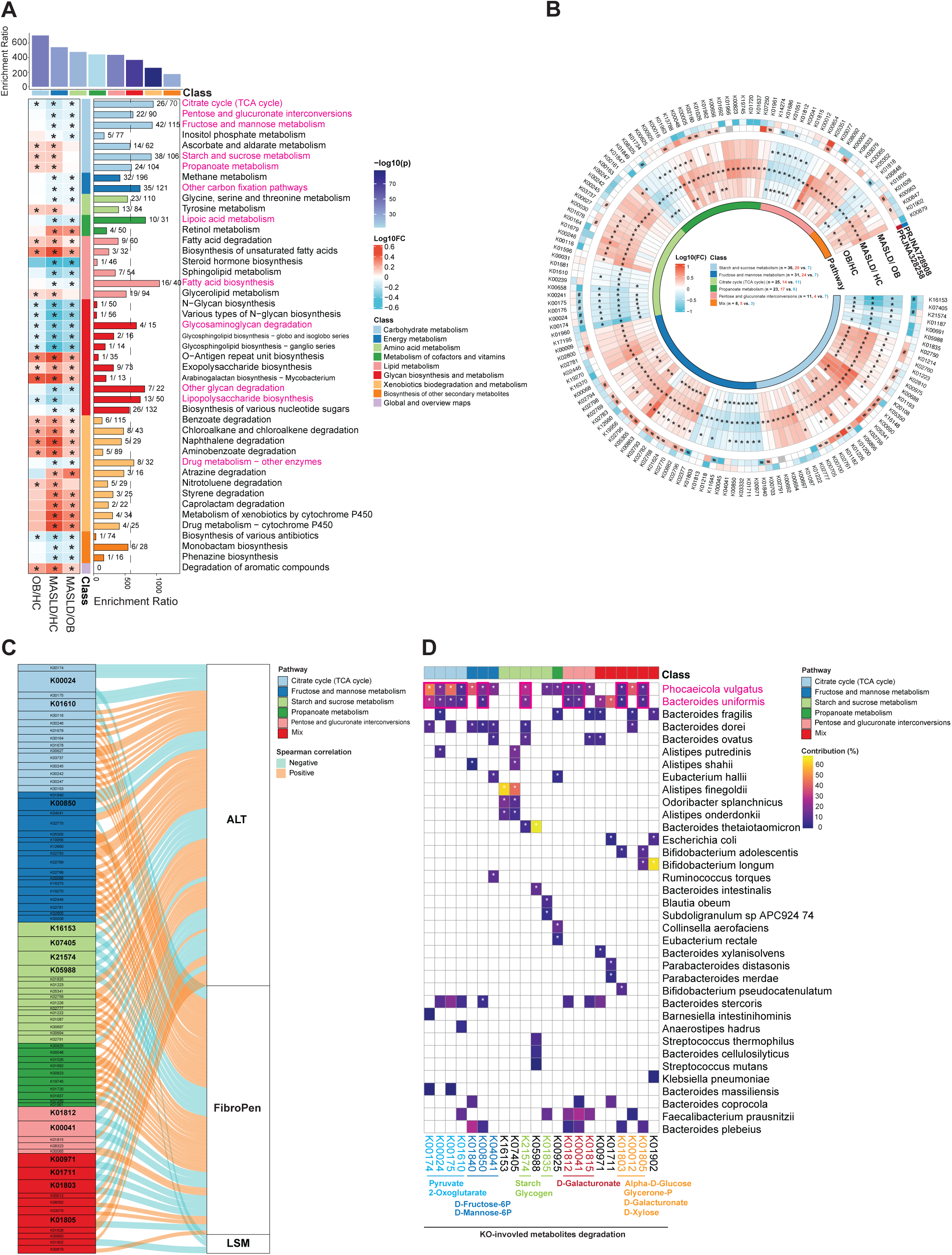
Distinctly Altered and Enriched Microbial Function. **(A)** Enrichment ratios of significantly altered pathways: the top panel shows the enrichment ratio for the class of the pathways, the bottom-left panel displays distinctly altered pathways, and the bottom-right panel presents their respective enrichment ratios. Pathways with an enrichment ratio greater than 600 are highlighted in red. **(B)** Log-transformed fold change values of significantly altered KOs (KEGG Orthology) in carbohydrate metabolism pathways with at least 10 significantly different KOs between HC, OB, and MASLD (inner panel). Fold change values of KOs in public cohorts (outer panel). The number of significantly altered KOs is summarized for each pathway. Increased KOs are shown in orange, and decreased KOs in blue. *FDR *P* < 0.05, # *P* < 0.05. **(C)** Correlation between significantly altered KOs in carbohydrate metabolism and clinical indices (ALT, Fibro-Pen, and LSM). The line thickness corresponds to the absolute value of the correlation coefficient. **(D)** The contribution (%) of the top 5 species to the KOs that negatively correlate with key clinical indices (ALT, Fibro-Pen, and LSM). The abundance of orthologous genes from species mapped to the KOs was compared between HC, OB, and MASLD. \**P* < 0.05.

Moreover, we found a total of 157 significantly abundant KOs (48% of the total) were predominantly mapped to carbohydrate metabolism pathways, with 105 KOs increased and 52 KOs decreased in MASLD (**Table S4**). Specifically, the pathway of starch and sucrose metabolism had the greatest number of differential KOs (n = 36). For example, K16153 and K21574 were progressively decreased from HC to OB to MASLD (FDR *P* < 0.05), and their changes were also validated in the public pediatric cohort (*P* < 0.05, **Figure 3B**). The next dysregulated pathway was fructose and mannose metabolism, despite an overall suppression (**Figure 3A**), there were 24 KOs increased in MASLD, including K02796, K00882, K01624, K02768, K02782 and K02793 showing a progressive increase from HC to OB to MASLD (FDR *P* < 0.05), with similar findings in the public pediatric cohort (*P* < 0.05). In contrast, there were 7 KOs decreased in fructose and mannose metabolism, especially K01840, K00971, K01711, K03332, K00850 and K04041, which progressively decreased from MASLD to OB to HC. Thirdly, there were 25 KOs significantly changed in the TCA cycle, with K00024, K00174, K00176, K00175 and K00241 showing a consistent decrease from MASLD to OB to HC (FDR *P* < 0.05), which were also confirmed in the public pediatric cohort. Taken together, these results suggest that gut dysbiosis in MASLD might lead to significant disruptions in carbohydrate metabolism, particularly in the starch and sucrose metabolism, fructose and mannose metabolism pathways, and TCA cycle.

### Carbohydrate Metabolism Disruption in MASLD Driven by the reduction of *B. uniformis* and *P. vulgatus*

To further investigate the relationship between significantly altered carbohydrate-associated KOs and disease severity (i.e., and ALT, Fibro-Pen, and LSM), we conducted Spearman’s correlation analysis and generalized linear regression, adjusting for potential confounders such as age, gender, and BMI z-score. After adjustment, the greatest number of significant correlations were observed in the TCA cycle, followed by fructose and mannose metabolism, as well as starch and sucrose metabolism (**Figure 3C**). Notably, K00024 in the TCA cycle showed significant negative correlations with ALT, Fibro-Pen, and LSM (*P* < 0.05). Additionally, K01610 in the TCA cycle, K00850 in the pathway of fructose and mannose metabolism, K16153, K07405, K21574, and K05988 in the pathway of starch and sucrose metabolism, and K01812 and K00041in the pathway of pentose and glucuronate interconversions exhibited significantly negative correlations with ALT and Fibro-Pen (*P* < 0.05). In contrast, K02770 and K02769 in fructose and mannose metabolism were positively correlated with ALT and Fibro-Pen (*P* < 0.05).

Next, we explored the role of gut microbiota in the regulation of carbohydrate metabolism by tracing the abundance of bacterial species-homologous genes within each KO. Interestingly, *P. vulgatus* and *B. uniformis* were found with the greatest contributions to those KOs that were negatively correlated with ALT, Fibro-Pen, and LSM (**Figure 3D**). Specifically, the decrease of *P. vulgatus* and *B. uniformis* contributed to the reduced expression of K00024, K00174, K00175, and K01610 in the TCA cycle, which were involved in the degradation of pyruvate and 2-oxoglutarate in MASLD. *P. vulgatus* and *B. uniformis* were also associated with the reduction of K01840, K00850, and K04041 in the fructose and mannose metabolism pathway, which were related to the degradation of D-fructose-6P and D-mannose-6P. In contrast, *Escherichia coli* was the species with the most significant contribution to KOs positively correlated with MASLD phenotypes, particularly those involved in the production of succinate, D-fructose-6P, D-glucose-6P, and D-fructose (**Figure S2**). Overall, the depletion of *B. uniformis* and *P. vulgatus* disrupted the microbial degradation of 2-oxoglutarate and monophosphate monosaccharide, while *E. coli* promoted their production, suggesting a potential mechanism in pediatric MASLD.

### Gut Microbiota-Encoded CAZyme Family Dysregulation in Pediatric MASLD

To specifically dissect the disruption of carbohydrate metabolism by gut microbiota in pediatric MASLD, we annotated CAZyme families using the CAZyme database. Six CAZyme categories were identified: glycoside hydrolases (GHs), carbohydrate esterase (CE), glycosyltransferases (GTs), polysaccharide lyases (PLs), carbohydrate-binding modules (CBMs), and auxiliary activities (AAs). In the MASLD group, GHs, CE, GTs, PLs, and CBMs were significantly reduced, while AAs levels were elevated (P < 0.05, **Figures 4A**).

**Figure 4.**
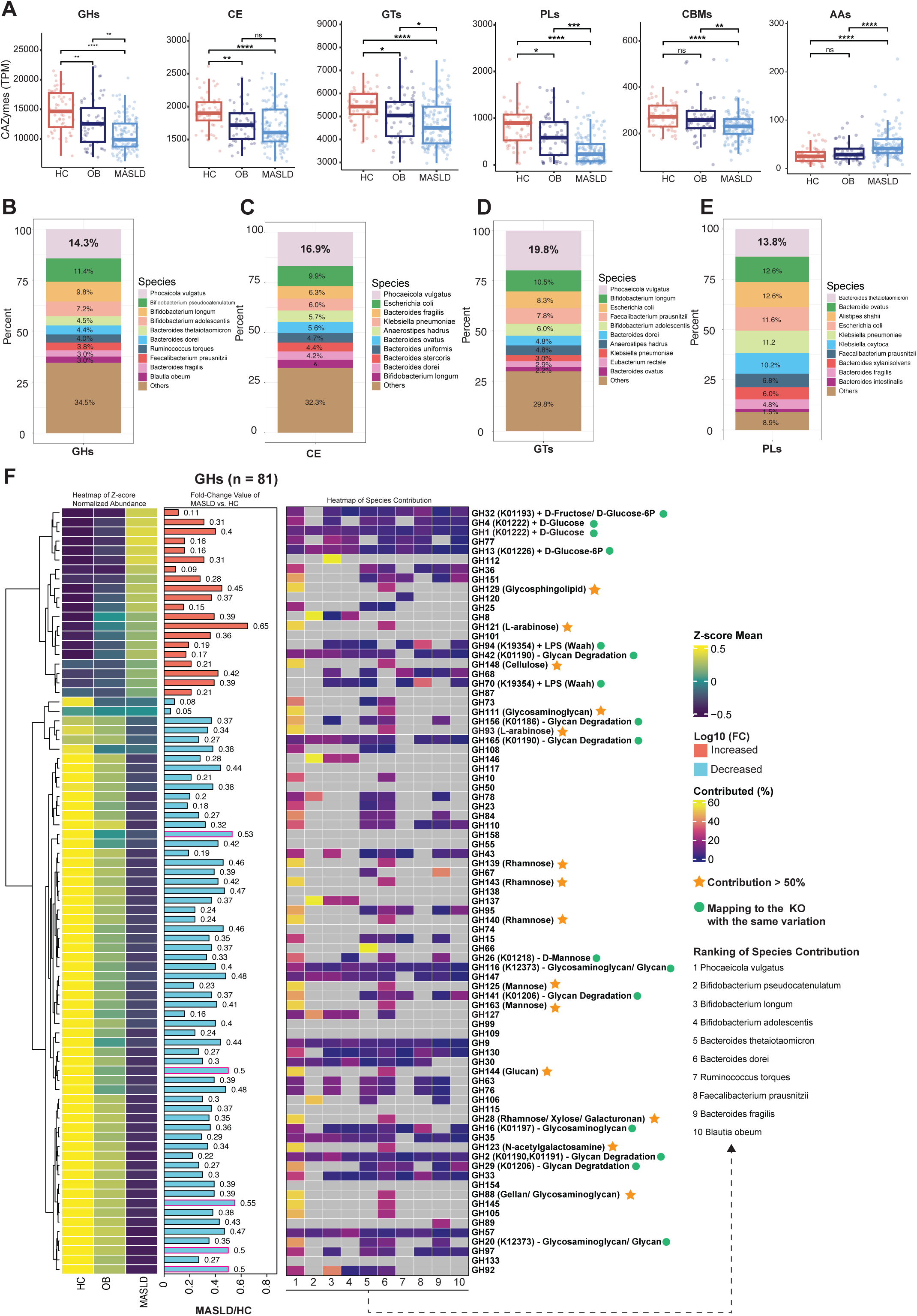
Dysregulation of Microbial CAZyme Family in Pediatric MASLD. **(A)** The CAZyme classes that significant difference between HC, OB and MASLD. GHs: glycoside hydrolases; CE: carbohydrate esterase; GTs, glycosyltransferases; PLs, polysaccharide lyases; CBMs, carbohydrate-binding modules; AAs, auxiliary activities. ****P < 0.0001, ***P < 0.001, **P < 0.01, *P < 0.05, ns denoted no significance. (**G-E**) Top 10 species contributing to each CAZyme class. (**F**) Z-score transformed mean of CAZyme families in GHs that were significantly different between HC, OB, and MASLD (first panel). Log-transformed fold change values between MASLD and HC (second panel). Fold change values less than 0 are presented as absolute values and are represented by blue bars, while values greater than 0 are shown as red bars. CAZyme families that significantly differ between simple and advanced MASLD are highlighted in red. The contribution (%) of the top 5 species to each GH family is displayed as a heatmap (third panel). The GH families that correspond to significantly different KEGG Orthologs (KOs) are highlighted with green circles. GH families with more than 50% contribution from *Phocaeicola vulgatus* are indicated by orange stars.

By tracing species-level contributions to KOs and matching EC numbers within each CAZyme family, we identified *P. vulgatus* as the primary contributor to GHs (14.3%), CE (16.9%), and GTs (19.8%), while *B. thetaiotaomicron* was mainly responsible for PLs (13.8%) (**Figure 4B-E**). Furthermore, we observed differential abundance of CAZyme families across HC, OB, and MASLD groups (FDR P < 0.05), including 81 GH families, 10 CE families, 20 GT families, 16 PL families, 11 CBM families, and 2 AA families (**Figure 4F** and **Figure S3**). Among the greatest number of significantly altered GH families, GH32 (mapping to K01193), GH4 (mapping to K01222), and GH1 (mapping to K01222) were notably increased, contributing to the production of D-fructose, D-glucose-6P, and D-glucose. Additionally, GH families with over 50% contribution from *P. vulgatus*, such as GH139, GH143, GH140, GH125, and GH163, were significantly decreased, which was associated with reduced rhamnose and mannose metabolism (**Figure 4F**). These findings suggest that *P. vulgatus* plays a crucial role in disrupting CAZyme families and impairing the metabolism of D-fructose, D-glucose, rhamnose, and mannose in children with MASLD, further validating the previous functional insights derived from the KEGG database.

### Integrated Metabolic and Microbial Interactions in MASLD Patients

Next, we conducted an untargeted metabolomics analysis on serum samples and identified 14 differentially abundant metabolites among three groups (HC, OB, and MASLD) involved in carbohydrate metabolism, including decreased levels of isocitrate, citrate, and malate in the TCA cycle, and increased levels of maltotriose, 2-oxoglutarate, pyruvate, glucose, and lactose (*P* < 0.05, **Figure S4A**). Metabolic enrichment analysis further revealed that TCA cycle and pentose and glucuronate interconversions were among top significantly enriched pathways in MASLD (FDR *P* < 0.05, **Figure 5A**). To gain a deeper understanding of host-microbiota interactions in energy and carbohydrate metabolism, we conducted a targeted metabolomics analysis on central carbon metabolism. A total of 65 metabolites with a prevalence greater than 60% of all the samples were selected for differential analysis. Among these, 64 were host-microbiota co-metabolized, while one originated exclusively from microbial metabolism (**Figure S4B**). We found 35 metabolites to be significantly different across HC, OB, and MASLD (*P* < 0.05, **Figure 5B**). Specifically, 17 metabolites (e.g., glycine, fumarate, maleic acid, cis-aconitate, and malate) were decreased, while 18 metabolites (e.g., D-glucose, D-fructose, N-acetylglucosamine 1-phosphate, adenosine-5’-diphosphate (ADP), inosine, and uridine 5’-monophosphate) were increased in MASLD. Consistent decreases in cis-aconitate and malate in the TCA cycle were observed across both untargeted and targeted metabolomics analyses. Correlation analysis showed that cis-aconitate, malate, fumarate, maleic acid, and glutamine were inversely associated with both ALT and Fibro-Pen, whereas D-glucose and D-fructose exhibited positive correlations with these markers (P < 0.05, **Figure S4D**). These results indicate that dysregulation of TCA cycle and monosaccharide metabolites may play a critical role in the development of pediatric MASLD.

**Figure 5.**
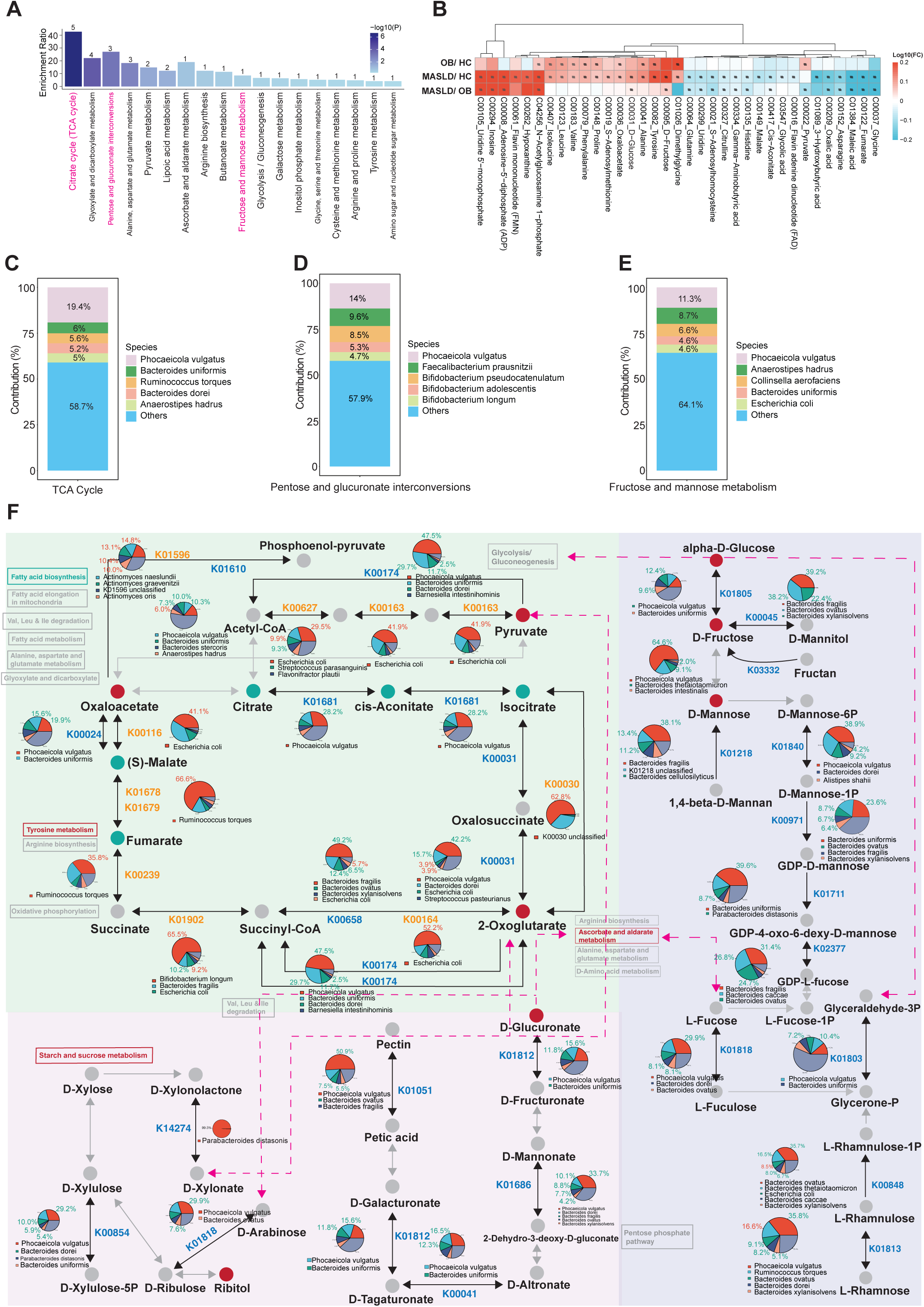
Integrated Metabolic Network of Host and Microbiota. **(A)** Enrichment ratio of each metabolic pathway, ranked by significance level. **(B)** Log-transformed fold change values of metabolites that were significantly different between HC, OB, and MASLD groups. **(C-E)** The contribution (%) of the top 5 species to the pathways of TCA cycle (C), pentose and glucuronate interconversions (D), and fructose and mannose metabolism (E). **(F)** Network diagram illustrating the TCA cycle, fructose and mannose metabolism, and pentose-glucuronate interconversion pathways, integrated with significantly changed KEGG Orthologs (KOs), species contributions to KOs, and metabolites. Significantly increased KOs are indicated in orange, while significantly decreased KOs are shown in blue. The contribution of the top 5 species to each KO is depicted in pie charts, with contributions showing significant differences between groups highlighted in red (increased in MASLD) and green (decreased in MASLD). Significantly different metabolites are represented as red circles (increased in MASLD) and green circles (decreased in MASLD).

We further summarized bacterial species encoded with key enzymes in the TCA cycle, pentose and glucuronate interconversions, and fructose and mannose metabolism. Consistent with CAZyme database analysis, *P. vulgatus* emerged as the primary contributor to these three key metabolic pathways, contributing 19.4%, 14%, and 11.3%, respectively (**Figure 5C-E**). By specifically integrating the differential KOs, contributing bacteria, and associated metabolites within these pathways, we outlined the dysregulated metabolic network underlying pediatric MASLD (**Figure 5G**). This network revealed that TCA cycle dysfunction was primarily associated with *P. vulgatus* and *B. uniformis*, leading to a reduced conversion of pyruvate to citrate, cis-aconitate, and isocitrate. Meanwhile, *E. coli* and *R. torques* dominated the increased metabolism of 2-oxoglutarate to succinate, fumarate, (S)-malate, and oxaloacetate (**Figure 5G**, top-left panel). The reduced pathway of fructose and mannose metabolism was primarily attributed to the decreased abundance of *P. vulgatus*, *B. uniformis*, and *B. thetaiotaomicron*, which might lead to diminished degradation of alpha-D-glucose, D-fructose, D-mannose, and L-rhamnose, resulting in significantly increased levels of alpha-D-glucose and D-fructose (**Figure 5G**, right panel). The reduction of pentose and glucuronic acid metabolism was linked to the reduced abundance of *P. vulgatus*, *B. uniformis*, and *P. distasonis*, which impaired the degradation of D-arabinose, D-xylose, and D-xylonate (**Figure 5G**, bottom-left panel).

In this disrupted metabolic network, pathways including pentose and glucuronate interconversions, fructose and mannose metabolism overlap with ascorbate metabolism, further influencing the TCA cycle. The level of ascorbate was significantly reduced in MASLD, as indicated by the untargeted metabolomics data (**Figure S4C**). These results suggest that the reduction of core symbiotic bacteria, such as *P. vulgatus*, *B. uniformis*, *P. distasonis*, and *B. thetaiotaomicron*, may affect multiple carbohydrate metabolism pathways, influencing monosaccharide metabolism, including D-glucose, D-fructose, D-mannose, L-rhamnose, and D-xylose. This disruption could lead to dysregulation of TCA energy metabolism, potentially contributing to the pathogenesis of pediatric MASLD.

### The Reduction of *P. distasonis*, and *B. uniformis* Contributed to The Gut microbial Dysbiosis and Imbalanced Carbohydrate Metabolism As Core Species in MASLD Progression

To further characterize microbial alterations in children with advanced MASLD who are at higher risk of MASH and fibrosis, we divided MASLD subjects into two subgroups: simple and advanced MASLD stratified by ALT, LSM, and FibroPen levels (**Figure S5A**). Advanced MASLD patients exhibited more pronounced dyslipidaemia and glucose metabolism disorders (**Table S5**). Overall, gut microbial diversity was comparable between subgroups (**Figure S5B**). We identified 34 species with differential abundance, among which *D. formicigenerans* was the only species consistently aligned with comparisons among HC, OB, and MASLD, showing further increase in advanced MASLD (**Figure S5C**, *P* < 0.05). *Coprococcus comes*, *Dysosmobacter sp. BX15*, and *Enterocloster aldenensis* exhibited consistent changes with public datasets. Co-abundance network analysis revealed persistent correlations between *P. vulgatus* and *B. uniformis*, and between *P. vulgatus* and *P. distasonis*, observed across networks derived from our advanced MASLD dataset, a public MASH dataset, and a public fibrosis dataset (**Figure S5D-E**, **Table S6**). Collectively, these results suggest a potential core dysbiotic guild constructed by *P. vulgatus*, *P. distasonis*, and *B. uniformis* may also play a crucial role in MASLD progression.

Overall, functional segregation was similar between subgroups (**Figure S5F**). Functionally differential analysis identified 97 differentially abundant KOs, with carbohydrate metabolism (n = 22) being the most affected (**Figure S5G**). Notably, K02796 in the fructose and mannose metabolism, and K01819 and K02082 in galactose metabolism, were consistently increased in advanced stages and in MASH of public datasets (**Figure S5G**, inner panel). Also, K05305 and K00882 showed further increases in the advanced stage, while K01805 further decreased. These changes, all related to fructose and mannose metabolism, were consistent with the results from the three major group comparisons (**Figure S5G**, outside panel), implicating a dysfunction in fructose and mannose metabolism during MASLD progression. Additionally, K01619, K00041 and K03738 in pentose and glucuronate interconversions and K01681, K00244 and K00240 in TCA cycle were found to be decreased in advanced MASLD. Species-level contributions to these metabolic shifts were further identified. *Clostridioides difficile* and *Eggerthella lenta* were found as the contributors to the decreased KOs K00001, K00127 and K00244 in carbohydrate metabolism. *P. vulgatus* and *B. uniformis* contributed to the decreased K01805, K01619 and K00041, which were involved in the fructose and pentose metabolism, despite their abundance remaining unchanged (**Figure S5H**). Conversely, *Blautia obeum* and *Blautia sp* showed significant enrichment and were associated with increased KOs in carbohydrate metabolism (**Figure S5I**). These findings suggest that *P. vulgatus* and *B. uniformis*, as part of the core dysbiotic guild, may drive changes in carbohydrate metabolism, potentially contributing to MASLD progression.

## Discussion

This study provides the cross-cohort characterization of gut microbial dysbiosis in pediatric MASLD, marked by a decline in core microbial commensals, including *P. vulgatus*, *B. uniformis*, *P. distasonis*, and *B. thetaiotaomicron*. These alterations primarily affect microbial functions related to carbohydrate metabolism. By integrating analyses of the gut microbiome, functional enzymes, and serum metabolites, we uncover a disrupted metabolic network driven by the reduction of these core gut symbionts and their coordinately dysregulated metabolites, including glucose, fructose, cis-aconitate, and malate, which might contribute to the development of pediatric MASLD.

The core microbiome, a set of microbial taxa with distinct genomic or functional traits specific to host or environment, plays a key role in shaping microbial structure and function [14, 16, 48]. Network analysis and co-occurrence patterns are increasingly used to identify these core taxa, which, regardless of their abundance, have a significant impact on the microbiome both individually and within microbial guilds [49, 50]. Recently, microbiologists have explored the diagnostic potential of microbial guild-level signatures to better understand their structural and functional relationships [14]. Our cross-sectional study identified marked depletion of species, such as *P. vulgatus*, *B. uniformis*, *P. distasonis*, and *B. thetaiotaomicron*, a pattern replicated in a public pediatric cohort from UK. Notably, *B. uniformis* and *B. thetaiotaomicron* showed significant correlations with MASLD markers (ALT and FibroPen) after adjustment of confounders age, sex and BMI z-score. These four species formed a robust co-occurrence guild validated across three independent public cohorts, with *P. vulgatus* and *B. uniformis* ranking among the top 10 most abundant species in the human gut. A previous study has recognized *Bacteroides* as a foundational genus, numerically dominant, and integral in shaping interactions [9]. Our findings further refine this understanding by demonstrating that *P. vulgatus* and *B. uniformis* act as both foundational and core species, with more pronounced co-abundance disturbances in children with obesity and MASLD compared to healthy controls. Therefore, the combined reduction of *P. vulgatus*, *B. uniformis*, *P. distasonis*, and *B. thetaiotaomicron* likely represents a core guild disruption, leading to structural imbalances in the gut microbiota of pediatric MASLD patients.

In addition to microbial structural disturbances, we observed metabolic perturbations in microbial functions, characterized by significant alterations in carbohydrate metabolism. Specifically, the highest enrichment ratios were observed in the decreased pathways of TCA cycle, pentose and glucuronate interconversions, and fructose and mannose metabolism. *P. vulgatus* and *B. uniformis,* as core guild constituents, were found to encode the KOs that were inversely associated with ALT, FibroPen, or LSM. These changes involved in carbohydrate metabolism including pyruvate catabolism, D-fructose-6P /D-mannose-6P processing, α-D-glucose utilization, and glycerone-P metabolism. Fructose phosphorylation generates D-fructose-6P, which is isomerized to D-glucose-6P for glycolytic ATP production, with pyruvate subsequently oxidized in the mitochondria via the TCA cycle [51, 52]. Gut has been regarded as a major source of circulating TCA cycle intermediates [53]. By integrating differential abundant functional pathways, KOs, and metabolites from both targeted and untargeted metabolomics, we provide a more detailed view of the disrupted metabolic network centered on the TCA cycle. Depletion of *P. vulgatus* and *B. uniformis* might affect the conversion of pyruvate to citrate, cis-aconitate, and isocitrate in the TCA cycle in MASLD patients. Notably, cis-aconitate, which showed a negative correlation with ALT in our data, is a precursor of itaconate with anti-inflammatory property [54, 55]. Thus, the reduced cis-aconitate levels in MASLD patients might exacerbate inflammatory responses.

Concurrently, *P. vulgatus*, *B. uniformis*, *P. distasonis*, and *B. thetaiotaomicron* were found to be the primary contributors of the KOs in the degradation of elevated α-D-glucose, D-fructose, and D-mannose to L-rhamnose. Elevated levels of α-D-glucose and D-fructose have been linked to increased inflammatory cytokine production (IL-1, IL-2, IL-6) in the intestines of mice [56], which affect the development of MASLD. Plasma mannose level was also significantly elevated in obese patients [57], while L-rhamnose has been shown to promote UCP1-dependent thermogenesis and improve obesity in mice [58]. Moreover, these species were found to encode the KOs associated with the metabolism of D-glucuronate, D-xylose, and D-arabinose in pentose and glucuronate interconversions. D-xylose has been documented for its therapeutic effects on NAFLD by suppressing macrophage-derived IL-1/IL-6 [59]. L-arabinose intervention in high-fat-diet mice was reported to enrich *Bacteroides* and promoted short-chain fatty acid production, thereby suppressing obesity [60]. The reduced pentose and glucuronate interconversion pathways, along with fructose and mannose metabolism, ultimately enter the 2-oxoglutarate metabolism in the TCA cycle through ascorbate and aldarate metabolism. Zheng et *al*. recently reported that elevated fructose/ascorbate ratio is positively associated with the increased serum uric acid level [61], strongly correlating with MASLD. Consistently, lower ascorbate levels were observed in our MASLD patients. Additionally, we identified *P. vulgatus* as a primary contributor to GH families through CAZyme database, which are involved in the metabolism of D-glucose, D-fructose, mannose, rhamnose, and L-arabinose. In advanced MASLD, the dysbiotic microbial network retained stable consisting of *P. vulgatus*, *B. uniformis*, *P. distasonis* as core guild species, which were further validated by independent cohorts. Taken together, our data suggest that the reduction of four core guild species, particularly *P. vulgatus*, may disrupt metabolic networks involved in monosaccharide metabolism and energy production through the TCA cycle in pediatric MASLD. These disruptions may contribute to impair carbohydrate metabolism and anti-inflammatory responses, thereby playing a critical role in the pathogenesis of MASLD in children.

Despite an overall suppression of the metabolic network consisting of TCA cycle, fructose and mannose metabolism, and pentose and gluronate interconversion pathways, specific KOs displayed significant upregulation that primarily encoded by *E. coli*. These species are primarily responsible for encoding KOs that promote the conversion of fumarate to malate and malate to oxaloacetate within the TCA cycle. The increased level of oxaloacetate observed in children with MASLD may suggest an enhanced generation of phosphoenolpyruvate for gluconeogenesis, which could disrupt metabolic homeostasis in pediatric MASLD. The increased abundance of *E. coli* has been commonly observed in metabolically unhealthy individuals and those with excessive weight gain [62, 63]. Ju et *al*. reported that *E. coli* colonization, associated with a decrease in *Bacteroides* abundance, resulted in significant body weight gain, colonic inflammation, liver adipogenesis, and impaired glucose tolerance [64]. In contrast, oral administration of *P. vulgatus* and *B. thetaiotaomicron* respectively in colitis mouse models showed anti-inflammatory effects against *E. coli* growth [65, 66], suggesting the competitive relationship between these core guild species and *E. coli* in the carbohydrate metabolism within microbial community. In addition, animal studies demonstrated that *P. vulgatus* [67, 68], *P. distasonis* [69, 70], and *B. thetaiotaomicron* [71] could potentially restore gut barrier function by reducing LPS production, thereby improving metabolic health. Thus, our findings highlight the potential therapeutic value of these core guild species, which might synergistically inhibit pathogenic bacteria growth, improve carbohydrate and energy metabolism, and enhance anti-inflammatory cytokine responses.

Finally, the limitations of this study should be acknowledged. First, the observational nature of our study means that evidence supporting a causal relationship between dysregulated carbohydrate metabolism driven by core guild species and the development of MASLD remains lacking. Therefore, the pathophysiological relevance and functional roles of gut microbiota-driven events in MASLD require further exploration and confirmation through studies that involve specific modification of gut species and associated animal models. Second, while we have adjusted for major confounders in our statistical analyses, we were unable to account for covariates such as diet and physical activity, which could potentially influence the results. Third, although we have tried to classify the MASH/fibrosis-prone population based on the non-invasive diagnostic biomarkers, we were unable to achieve precise pathological subgrouping as compared to liver biopsy.

## Conclusion

Through an integrated dual-omics analysis, our study delineates gut microbial dysbiosis in pediatric MASLD with cross-cohort validation, characterized by the depletion of core species, including *P. vulgatus*, *B. uniformis*, *P. distasonis*, and *B. thetaiotaomicron*. These species form a core guild in MASLD, primarily influencing carbohydrate metabolism and contributing to a disrupted metabolic network involving key substrates such as glucose, fructose, and intermediates of the TCA cycle. Our findings suggest that the interplay between these core species and their associated metabolites may drive metabolic disturbances and inflammatory responses, playing a pivotal role in the pathogenesis of pediatric MASLD.

## Author contribution statement

**Jiating Huang**: Methodology, Data Curation, Formal analysis, Visualization, Writing-Original draft preparation, Writing - Review & Editing. **Xuelian Zhou**: Methodology, Investigation, Data Curation; **Huiying Wang**: Investigation, Data Curation. **Ana Liu**: Methodology, Investigation, Data Curation; **Junfen Fu**: Methodology, Investigation, Data Curation; **Guanping Dong**: Methodology, Investigation, Data Curation; **Ying Shen**: Methodology, Investigation, Data Curation; **Wenqing Xiang**: Methodology, Investigation, Data Curation; **Jeffrey Schwimmer**: Methodology, Investigation. **Gang Yu**: Methodology, Investigation, Data Curation. **Jian Huang**: Methodology, Investigation. **Yingping Xiao**: Methodology, Investigation. **Yan Ni**: Supervision, Project administration, Funding acquisition, Conceptualization, Methodology, Writing-Original draft preparation, Writing - Review & Editing.

