## Supplemental Figure S1-S5 for "Integrated Metagenomics and Metabolomics Studies Reveal Core Bacterial Guild Regulating Carbohydrate Metabolism in Pediatric MASLD"

**Supplementary Figure 1**

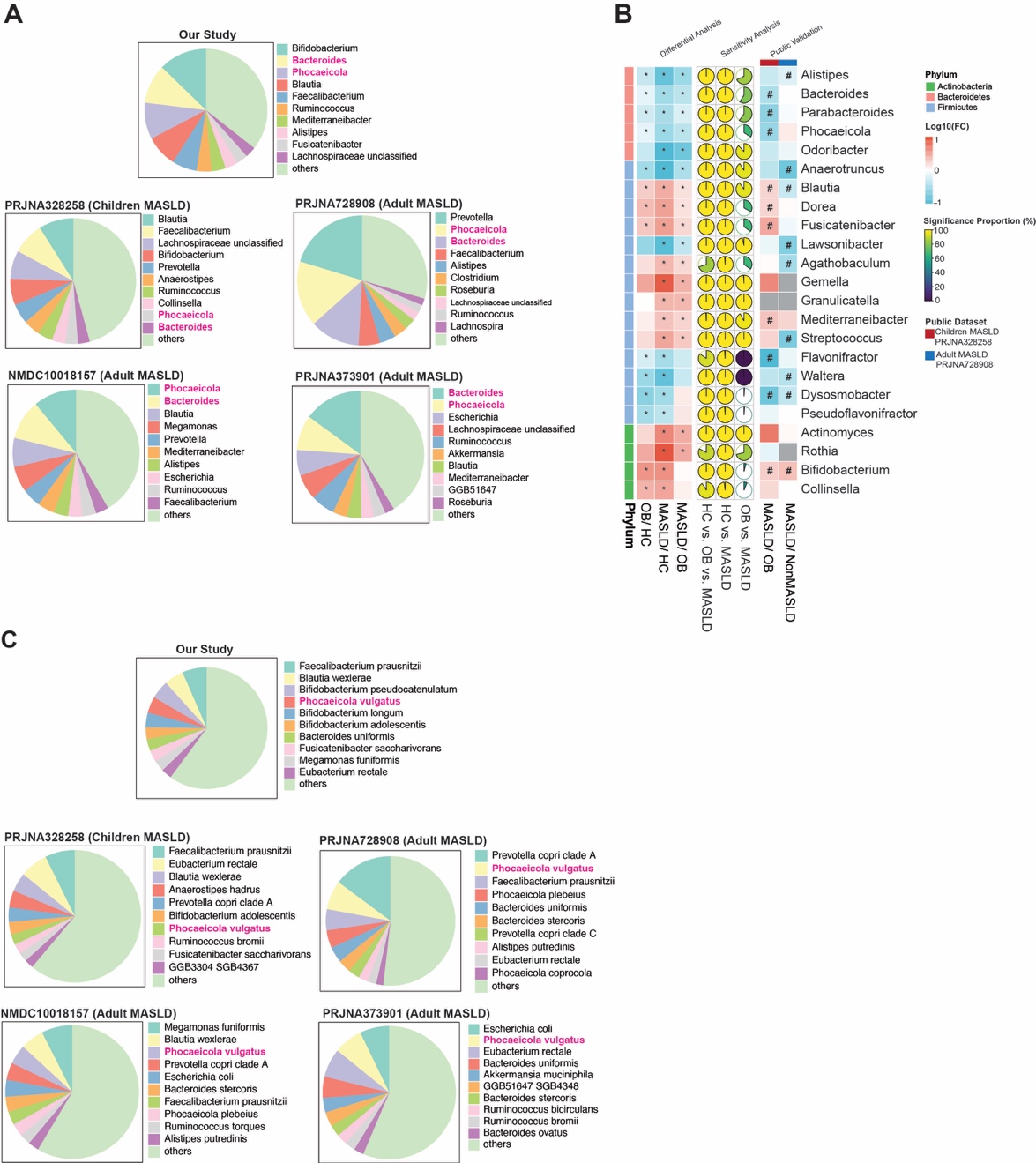

**Figure. S1. Microbial variation in pediatric MASLD, related to Figure 1.**

**(A)** Ranking of the top 10 most abundant microbial genera in each dataset, with genera shared across all datasets highlighted in red. **(B)** Log-transformed fold change values of microbial genera with significant variations (left panel), sensitivity analysis (middle panel), and validation using public datasets (right panel). Genera were compared across the three groups using the Kruskal-Wallis test, with genera having FDR-adjusted *P* < 0.05 selected for pairwise comparison via the Wilcoxon rank sum test. *FDR *P* < 0.05, #*P* < 0.05. **(C)** Ranking of the top 10 most abundant microbial species in each dataset, with genera shared across all datasets highlighted in red.

**Supplementary Figure 2**

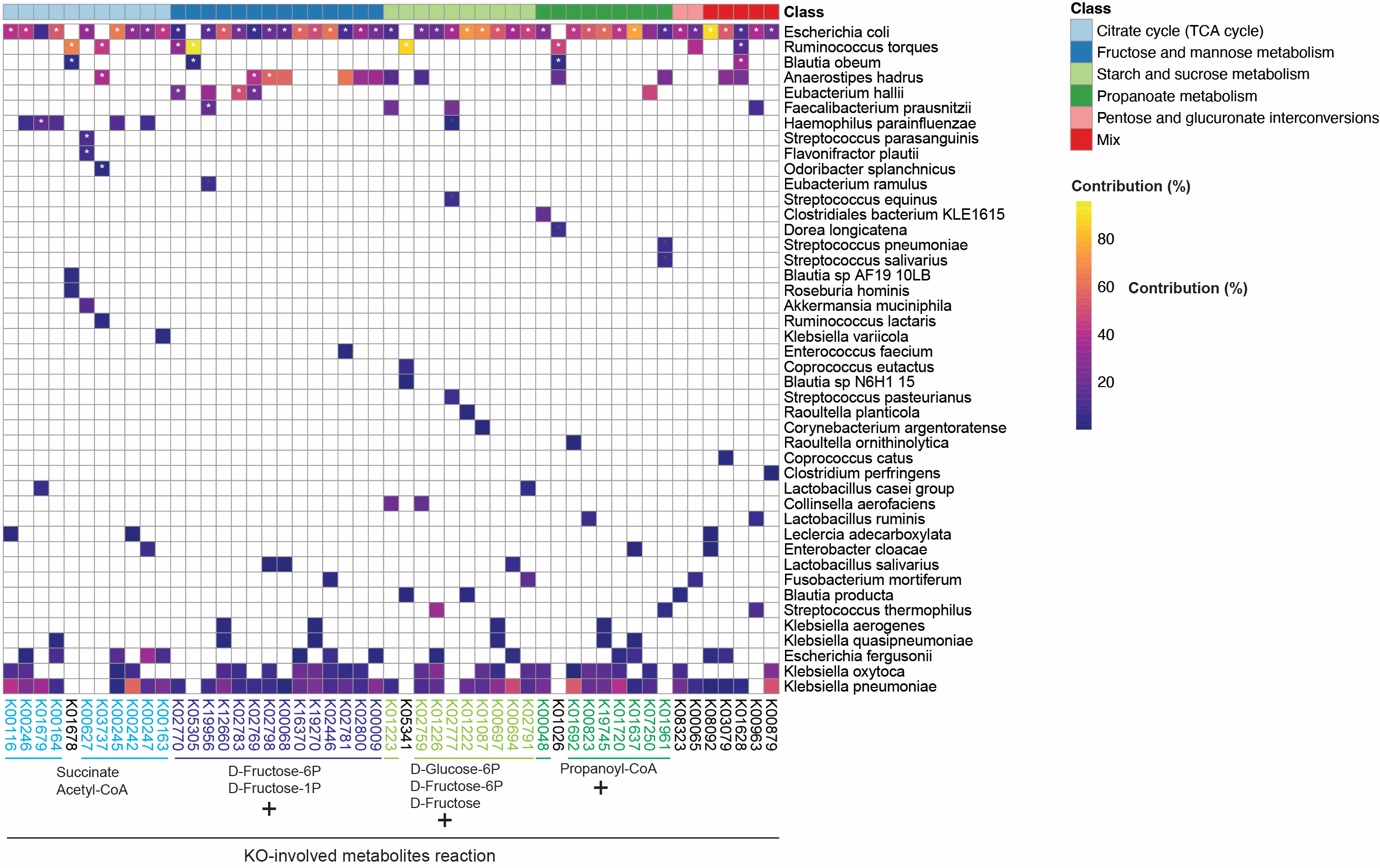

**Figure S2. The species contribution heatmap of the KOs significantly correlated with ALT, Fibro-Pen, and LSM, related to Figure 3C-D.**

The contribution (%) of the top 5 species to KOs negatively correlated with key clinical indices (ALT, Fibro-Pen, and LSM). The abundance of orthologous genes from species mapped to the KOs was compared between HC, OB, and MASLD. **P* < 0.05.

**Supplementary Figure 3**

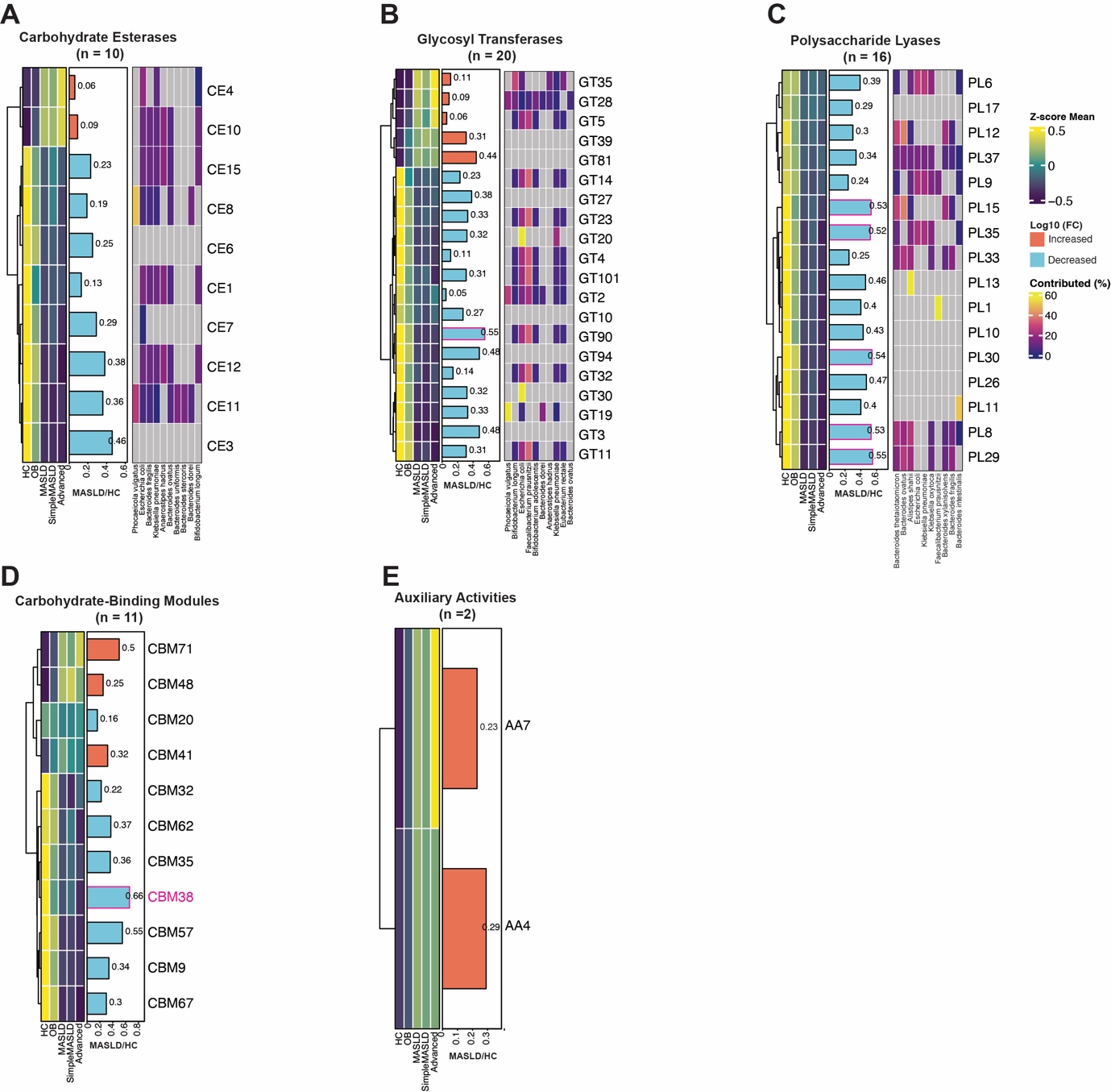

**Figure S3. The significant variation of CAZyme family in each class, related to Figure 4.**

**(A-E)** Z-score transformed mean of CAZyme families (first panel) in carbohydrate esterase (CE), glycosyl transferases (GT), polysaccharide lyases (PL), carbohydrate-binding modules (CBM) and auxiliary activities (AA). Log-transformed fold change values between MASLD and HC (second panel). Fold change values less than 0 are presented as absolute values and are represented by blue bars, while values greater than 0 are shown as red bars. CAZyme families that significantly differ between simple and advanced MASLD are highlighted in red. The contribution (%) of the top 5 species to each CE, GT and PL family is displayed as a heatmap (third panel).

**Supplementary Figure 4**

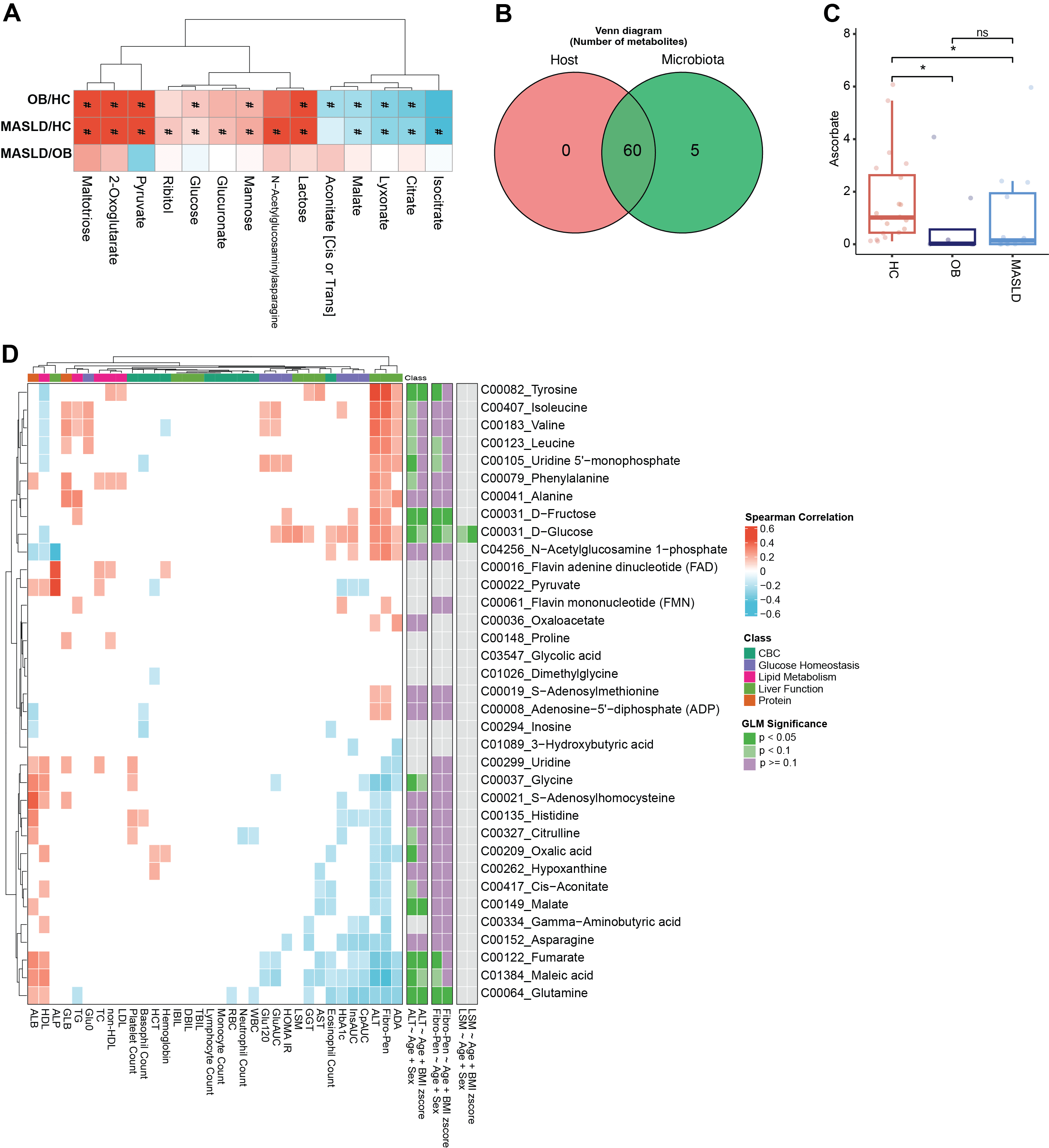

**Figure S4. The significant variation of metabolites and correlation analysis, related to Figure 5.**

**(A)** The concentration of D-glucose with significant differences between simple MASLD and advanced MASLD. ***P* < 0.01. **(B)** Venn plot of metabolites source. **(C)** The relative abundance of ascorbate from untargeted metabolomics, with significant variation between HC, OB, and MASLD. **P* < 0.05, ns indicates no significant difference. **(D)** Significant correlations (*P* < 0.05, Spearman's correlation) are represented in colour: red indicates positive correlation, while blue indicates negative correlation. The correlations between species and key clinical indices, including ALT, Fibro-Pen, and LSM, were further adjusted for confounders using generalized linear regression. After adjustment, correlations with *P* < 0.05 are shown in deep green, *P* < 0.1 in light green, and *P* ≥ 0.1 in purple.

**Supplementary Figure 5**

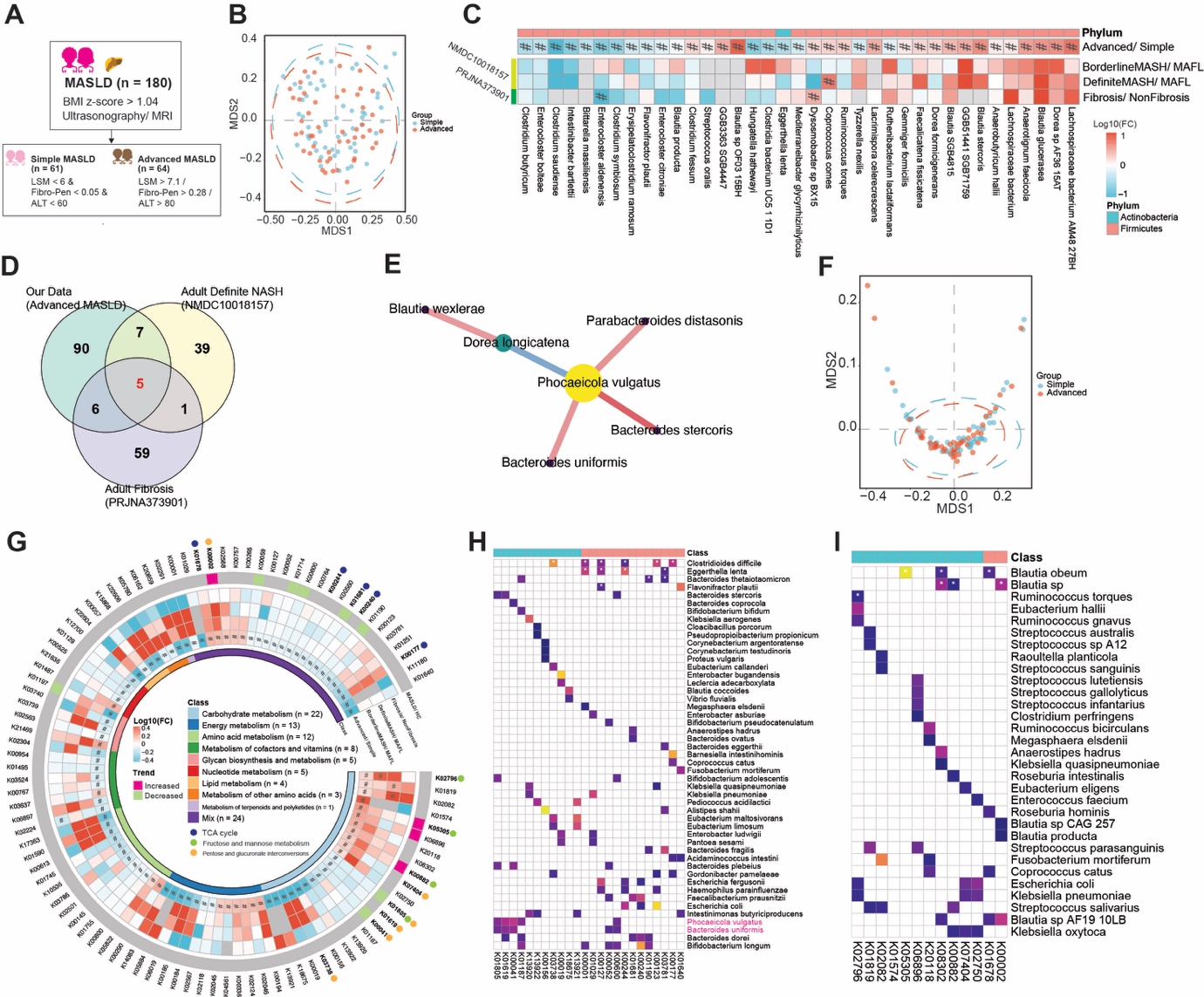

**Figure S5. Comparison between simple and advanced MASLD status.**

**(A)** Study design of MASLD subgroups. **(B)** Multidimensional scaling analysis of the taxonomic variation. **(C)** Log-transformed fold change values of microbial species with significant differences between simple and advanced MASLD status. #*P* < 0.05. **(D)** Venn diagram showing shared connections in advanced MASLD-associated networks. **(E)** Consistent correlations across advanced MASLD-associated networks. (**F**) Multidimensional scaling analysis of the functional variation. **(G)** Log-transformed fold change values of significantly altered KOs between simple and advanced groups (inner panel). Cross-validation of these changes with public datasets (inner panel) and consistent changes when comparing MASLD with HC. **(H-I)** Contribution (%) of the top 5 species to significantly altered KOs between simple and advanced groups (H: Decreased KOs; I: Increased KOs). **P* < 0.05.
